# Separation and quantification of 2-Keto-3-deoxy-gluconate (KDG) a major metabolite in pectin and alginate degradation pathways

**DOI:** 10.1101/2020.07.24.220400

**Authors:** Shiny Martis B, Michel Droux, Florelle Deboudard, William Nasser, Sam Meyer, Sylvie Reverchon

## Abstract

A rapid and sensitive High Performance Liquid Chromatography (HPLC) method with photometric and fluorescence detection is developed for routine analysis of 2-Keto-3-deoxy-gluconate (KDG), a catabolite product of pectin and alginate. These polysaccharides are primary-based compounds for biofuel production and for generation of high-value-added products. HPLC is performed, after derivatization of the 2-oxo-acid groups of the metabolite with o-phenylenediamine (oPD), using a linear gradient of trifluoroacetic acid and acetonitrile. Quantification is accomplished with an internal standard method. The gradient is optimized to distinguish KDG from its close structural analogues such as 5-keto-4-deoxyuronate (DKI) and 2,5-diketo-3-deoxygluconate (DKII). The proposed method is simple, highly sensitive and accurate for time course analysis of pectin or alginate degradation.

**Highlights:** A fluorescent based-HPLC method report the quantification of KDG, a metabolite originating from alginate and from pectin degradation pathways, using derivatization with o-phenylenediamine (oPD)

## Introduction

The major obstacle to use plant biomass for the production of fuels, chemicals, and bio-products, is our current lack of knowledge of how to effectively deconstruct cell-wall polymers for their subsequent use as feed stocks. Pectin plays an important role in biomass recalcitrance [1]. Pectins are complex branched polysaccharides containing high amounts of partly methyl-esterified galacturonic acid and low amounts of rhamnose, and carry arabinose and galactose as major neutral sugars. Due to their structural complexity, they are modifiable by many different enzymes, including polygalacturonases, pectate lyases, and esterases. Pectin-degrading enzymes and -modifying enzymes may be used in a wide variety of applications to modulate pectin properties or to produce pectin derivatives for food industry [2], as well as for pharmaceutical industry. Both, pectin and pectin-derived oligosaccharides act as anti-metastatic [3], cholesterol inhibiting [4], prebiotic and probiotic encapsulants [5]. In addition, pectin is widely used as a component of bio-nano-packaging [6,7].

In this context, numerous plant pathogenic microorganisms such as soft-rot bacteria (*Dickeya* species and *Pectobacterium* species) or necrotic fungi (*Aspergillus species*, *Botrytis species*) are effective in pectin degradation and represent a large resource of pectinases that generated the KDG metabolite [8–10]. In addition, novel metabolic pathways for hexuronate utilization in various proteobacteria have been recently described and all converge to the KDG metabolite formation [11,12].

Similarly, algae are a large group of marine vegetation that also emerge as feedstock for biofuel production [13,14]. Alginate is a unique structural polysaccharide in algae, abundant in the cell wall ensuring resistance to mechanical stress. It is a linear copolymer of α-L-guluronate and its C5 epimer β-D-mannuronate [15]. Alginate degrading enzymes include different types of alginate lyases generating various alginate oligosaccharides that are further converted to KDG. Alginate lyases are present in a wide range of marine and terrestrial bacteria [16]. In addition to biofuel production, alginate derivatives can be used as therapeutic agents such as anticoagulants [17] and tumor inhibitors [18]. Furthermore, alginate lyase also shows great potential applications in treatments of cystic fibrosis by degrading the polysaccharide biofilm of bacteria [19]. KdgF was identified as the missing link in the microbial metabolism of uronate sugars from pectin and alginate|20] (Figure 1).

**Figure 1:**
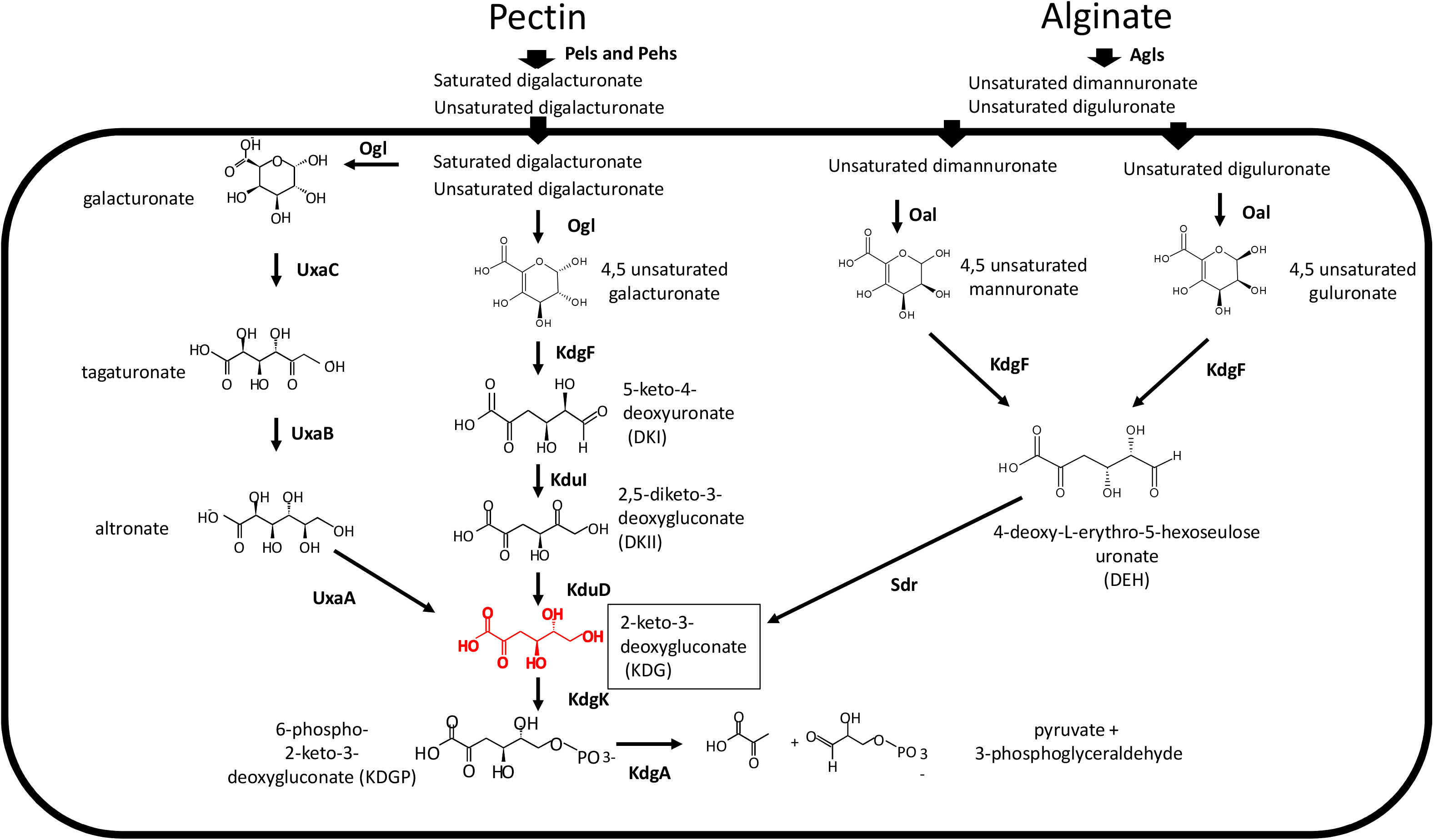
Schematic metabolic pathways leading to KDG (2-Keto-3-deoxy-gluconate) production from pectin and alginate degradation. Pectin degradation involved pectate lyases and polygalacturonases that cleaved the polygalacturonate, the demethylated pectin, into unsaturated and saturated digalacturonate, respectively. Inside the bacteria, these compounds are converted by Ogl enzyme into galacturonate and unsaturated galacturonate (for saturated digalacturonate) and into two molecules of unsaturated galacturonate (for unsaturated digalacturonate). KdgF converts unsaturated galacturonate into DKI. kduI enzyme then converts DKI to DKII and kduD subsequently converts DKII to KDG, 2-Keto-3-deoxy-gluconate. This compound is then phosphorylated by a specific kinase kdgK and finally cleaves by KdgA to yield pyruvate and phospho-glycerate. The galacturonate is degraded to KDG by the enzymes UxaC, UxaB, UxaA in the hexuronate pathway. Alginate degradation involved alginate lyases that cleaved the corresponding polymer into unsaturated dimannuronate and unsaturated diguluronate. Inside the bacteria, these compounds are converted by oligouronide lyase Oal into unsaturated mannuronate and unsaturated guluronate monomers. These 4,5-unsaturated monouronates are converted to DEH (4-deoxy-L-erythro-5-hexoseulose uronate) by the enzyme KdgF. DEH is then reduced into KDG by the Sdr enzyme. Enzymes stand for: Pels, pectate lyases; Pehs, polygalacturonases; Ogl, oligogalacturonate lyase; KdgF, responsible for linearisation of 4,5-unsaturated monouronates to linear ketonized forms; kduI, 4-deoxy-L-threo-5-hexosulose-uronate ketol-isomerase; kduD, 2-dehydro-3-deoxy-D-gluconate 5-dehydrogenase; kdgK, 2-dehydro-3-deoxygluconokinase; KdgA, 2-dehydro-3-deoxy-phosphogluconate aldolase; UxaC, D-galacturonate isomerase; UxaB, Altronate oxidoreductase; UxaA, Altronate dehydratase; Agls, alginate lyases; Oal, oligouronide lyase; Sdr, 4-deoxy-L-erythro-5-hexoseulose uronate reductase.

Therefore, quantification of KDG is of wide interest to analyze the metabolic flux of hexuronate, pectin and alginate catabolism from numerous microorganisms.

In this paper, we took advantage of our expertise on the plant pathogenic bacteria *Dickeya dadantii* and the availability of both chemical and genetic resources in order to develop methods for monitoring the breakdown of pectic polymers into monomers of interest. This bacterium attacks a wide range of plant species, including many crops of economic importance. Soft rot, the visible symptom, is mainly due to the degradation of pectin by pectate lyase (Pel) activity generating unsaturated oligogalacturonides [21,22]. *Dickeya* also produces low amounts of polygalacturonases (Pehs) generating saturated oligogalacturonides [23,24]. *D. dadantii* can utilize pectin as its sole carbon and energy source (Figure 1) [21]. The catabolic pathway of pectin (pectinolysis) is characterized at the genetic and biochemical level, showing that KDG, a catabolite product of pectin, plays a key role in pectinolysis by inactivating KdgR, the main repressor of this catabolic pathway [25–27]. The positive feedback loop between the extracellular pectin degradation by Pels and Pehs and the intracellular inactivation of KdgR by the metabolite KDG has been modelled [7,28]. Refining this type of model required a precise determination of the concentration of KDG present in the bacteria's growth environment. The conventional technique to determine the concentration of KDG so far is thiobarbituric assay [29]. Usually, detection of KDG is performed by cellulose thin-layer chromatography and spraying with either periodate-thiobarbituric acid or o-phenylenediamine [30]. This method is still in use to follow the activity of enzymes such as Aldo-Keto Reductase reducing alginate-derived 4-deoxy-L-*erythro*-5-hexoseulose uronic acid to KDG [31,32]. Recently, HPLC based UV-detection coupled to NMR was reported for identification of KDG when studying the purified 2-keto-3-deoxygluconate aldolase from *Sulfolobus solfataricus* [33,34].

These methods are not very accurate for biological application with cell crude extracts and culture media since they are not very specific to KDG alone, but rather quantify DKI, DKII, KDG and other periodate-oxidizing sugars or o-phenylenediamine-reacting compounds indistinctly. Here, we present a sensitive method to quantify KDG after derivatization with oPD and separation using HPLC coupled to fluorescence detection. The reaction of oPD with 2-oxoacids has been used in HPLC chromatographic determination when working with human urine and plasma in the yearly’s 1980 [35–37].

The technique reported here allows us to quantify KDG in complex mixture with precision and specificity with respect to functionally related compounds, as required for further regulatory models.

## Materials and Methods

### Chemicals

KDG (95%), oPD (o-phenylenediamine), HPLC grade Trifluoroacetic acid (TFA), periodic acid (HIO_4_), sodium arsenite (AsNaO_2_), thiobarbituric acid (TBA) were from Sigma Aldrich (France). Partially purified DKI and DKII (90%) was a gift by Guy Condemine. Optima LC-MS acetonitrile (ACN) was from Fisher UK. Polygalacturonate dipecta (PGA) was from Agdia laboratory (France). All HPLC eluates, standards and assays were prepared using MilliQ pure water with a specific resistance of at least 18 mΩ.

### Bacterial strains and culture conditions

All the experiments were performed with the wild type *Dickeya dadantii* (strain 3937) and its derived *kduI*, *kduD* and *kdgK* mutants [21, 38–41] (Table 1). Bacteria were grown at 30°C in M63 minimal medium supplemented with a carbon source: glucose (0.2% w/v) alone or in addition with PGA (0.4% w/v). Cells were collected in the late exponential growth phase when the bacterial population reached its maximum at 10-hours. The bacterial cells were separated from residual polygalacturonate by centrifugation.

**Table 1:** *Dickeya dadantii* strains used in the study

| Strain | Genotype | Reference |
| --- | --- | --- |
| 3937 | Wild Type | [31] |
| A2395 | *kduI::kan* | [32] |
| A797 | *kduD::kan* | [33] |
| A3999 | *kdgK::kan* | [34] |

### Sample collection

Bacteria were separated from the medium by centrifugation at 5000 g for 8 minutes. Cells and cultures medium samples were freeze dried with liquid nitrogen and stored separately at - 80°C. Bacterial pellet was resuspended with a 10 ml cold solution of 2.5 % (v/v) TFA, 50% (v/v) acetonitrile in MilliQ water. The dissolved lysed pellet was maintained at - 80°C for 15 minutes and thawed at room temperature. The solution was then centrifuged at 20000 g for 5 minutes. The clear supernatant containing the metabolites (10 ml) was separated from the pellet and then lyophilized overnight using a Christ 2,4 LD plus equipped with a trap before the vacuum pump (Thermofisher, France). The dry powder was dissolved in 1 ml of 0.1 M HCl and centrifuged again to remove particles. Culture medium (10 ml) was lyophilized and re-suspended in 1 ml 0.1 M HCl.

### Sample derivatization

200 μL of sample was incubated for 60 minutes at 60°C with 20 μL solution of 100 mM oPD prepared in 0.1 M HCl with agitation in an Eppendorf thermomixer as described previously [42,43], (Figure 2). Samples turn yellowish with a typical UV-visible spectrum of a quinoxaline derivative [44]. Appropriate amounts of the metabolites-adduct were then injected into HPLC for analysis. These conditions were optimized with respect to temperature, incubation time and concentration of oPD.

**Figure 2:**
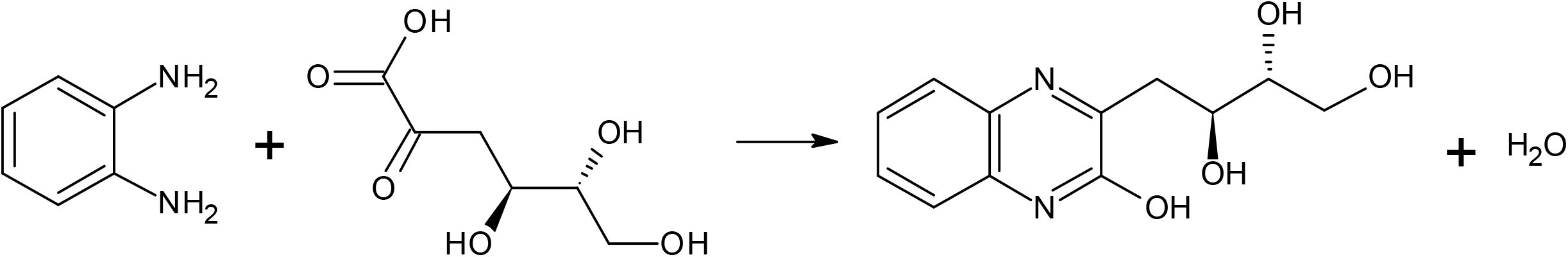
Derivatization of the α-diketones group of KDG with o-phenyldiamine to form quinoxaline derivative (oPD adduct).

### HPLC system and gradient elution

The HPLC system consisted of an Alliance HT Waters 2795 separation module and detection was performed with a Waters 2996 photodiode array detector interconnected with a fluorimeter SFM25 (Kontron). The oPD-adducts to be determined were passed through a 4.6×250-mm Uptisphere 5 μM HDOC18 column (Interchim) at 35°C and a flow rate of 1 mL/min. The elution protocol employed linear gradient as follow: initial 0 to 5 min, 100% A; 5 to 12 min, 75% A, 25% B; 12 to 20 min, 75% A, 25% B; 20 min to 30 min 75% A, 25% B to 100 % B; 30 min to 35 min, 100% B, then, 35 min to 37 min 100% B to 100% A and re-equilibration for 3 min in 100% A before a new run. Solvent A consisted of 0.005% trifluoroacetic acid (TFA) and solvent B contained 60% acetonitrile both in MilliQ water. The oPD adduct(s) was detected using a photodiode array detector (Waters 2996) set up at 338 nm, and with a SFM25 fluorimeter (Kontron) equipped with a 8 μL flow cell. Emitted fluorescence of the oPD adduct(s) was measured with an excitation wavelength of 338 nm and an emission wavelength of 414 nm (slit sizes, 15 and 10 nm respectively, and lamp power of 400 W). Quantification of metabolites was obtained by measuring peak areas using the chromatography data system Mass-Link software and assigned intensity to the peak (Waters).

### Thiobarbituric acid assay

A standard curve with increasing KDG concentration was generated using the procedure described [29,30]. Briefly, each amount of KDG in water (0.2 ml) was oxidized in presence of 0.25 ml of 25 mM HIO_4_ dissolved in 0.125 N H_2_SO_4_ for 20 min at room temperature, then 0.5 ml AsNaO_2_ 2% in 0.5N HCl was added to stop the reaction with shaking. Then, 2 ml of thiobarbituric acid 0.3% was added and the mixture was heated for 10 min at 100°C. After cooling, the optical density was measured at 510 nm in a spectrophotometer.

### Pectate lyase (Pel) activity

Assay of pectate lyase was performed on supernatants of bacterial cultures or on toluenized cell extracts as described previously [28]. Pectate lyase activity was determined by the degradation of PGA to unsaturated products that absorb at 230 nm. A molar extinction coefficient for unsaturated oligogalacturonides of 5200 was used. Pel activity was expressed as μmol of unsaturated products liberated per min and per ml of enzymatic extract, using the standard assay mixture described |28]. Specific activity is expressed as μmol of unsaturated products liberated min^−1^.mg^−1^ (dry weight) bacteria. Bacterial concentration was estimated by measuring turbidity at 600 nm, given that an optical density at 600 nm (OD_600_) of 1.0 corresponds to 10^9^ bacteria.mL^−1^ and to 0.47 mg of bacteria (dry weight) mL^−1^ [45].

## Results and discussion

The derivatization and subsequent HPLC procedure described here represents a successful improvement over the thiobarbituric method developed previously [24]. We used oPD as a probe for the ketone group of metabolites such as KDG, DKI and DKII (Figure 2). Figure 3 shows a representative chromatogram of KDG standard solution. The standard KDG has a retention time of around 16 minutes. From the estimated area of the respective peaks, the fluorescence intensity is approximately 2500-fold more sensitive than the UV absorbance intensity. Thus, fluorescence is more accurate for the resolution and quantitation of the metabolite of interest. The oPD adduct of the ketone group of KDG is characterized by a typical spectrum with a maximum absorption in the limit of the visible range at 338 nm [44]. Upon excitation at this wavelength, a maximum emission is measured at 414 nm (Figure 4A and B).

**Figure 3:**
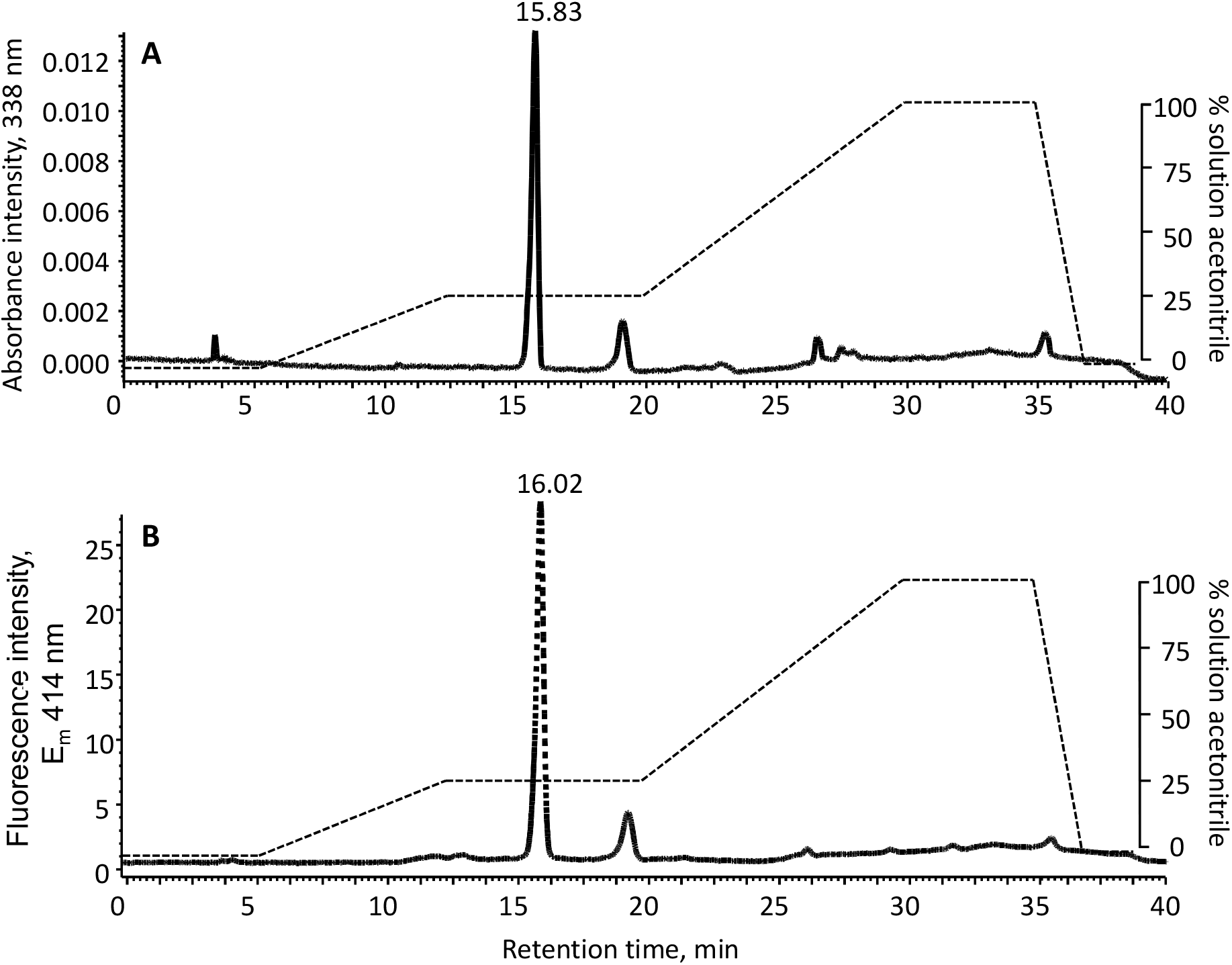
Chromatogram for the detection of 1 nanomole of standard KDG-oPD molecule adduct. A) Absorbance and B) Fluorescence detection.

**Figure 4:**
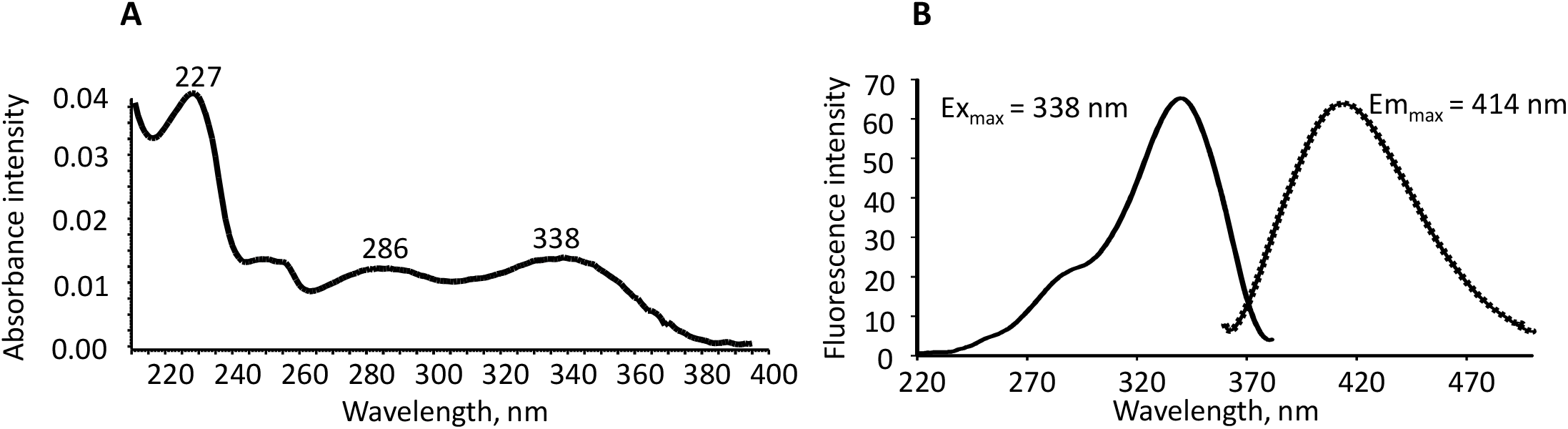
Physical properties of the KDG-oPD adduct. A) Characteristic of the absorbance spectrum of 1 nanomole of the KDG-oPD molecule adduct derived from the peak previously isolated by HPLC. B) Excitation and emission peaks of the purified KDG-oPD molecule adduct purified above. Spectra were performed in the elution buffer using a SFM25 spectrofluorimeter (Kontron).

The unsaturated digalacturonate pathway has DKI and DKII as intermediate metabolites before the formation of KDG (Figure 1). The properties of derivatization based on the presence of a ketone group led us to hypothesize that these metabolites would also be derivatized in our experimental conditions. We performed the same analysis starting from a partially pure DKI/DKII mixture (obtained by purification from *kduI/kduD* mutants, G. Condemine, personal communication). The closeness of the structures of DKI and DKII makes it difficult to differentiate them. Consistently, the corresponding chromatogram reveals a major single peak with a retention time of about 21 minutes for these compounds (Figure 5). We hypothesized that the signature spectrum of the KDG molecule would be comparable to that of DKI or DKII since the molecular structures are closely similar (see Figure 1). Only the predominant peak present in the absorbance chromatogram (Figure 5A) shows the typical spectrum signature of KDG as described in Figure 4A. We therefore optimized the gradient to discriminate the KDG from other similar structured metabolites produced in the media culture like DKI and DKII, allowing quantification from the intensity of a single specific peak (Figure 5).

**Figure 5:**
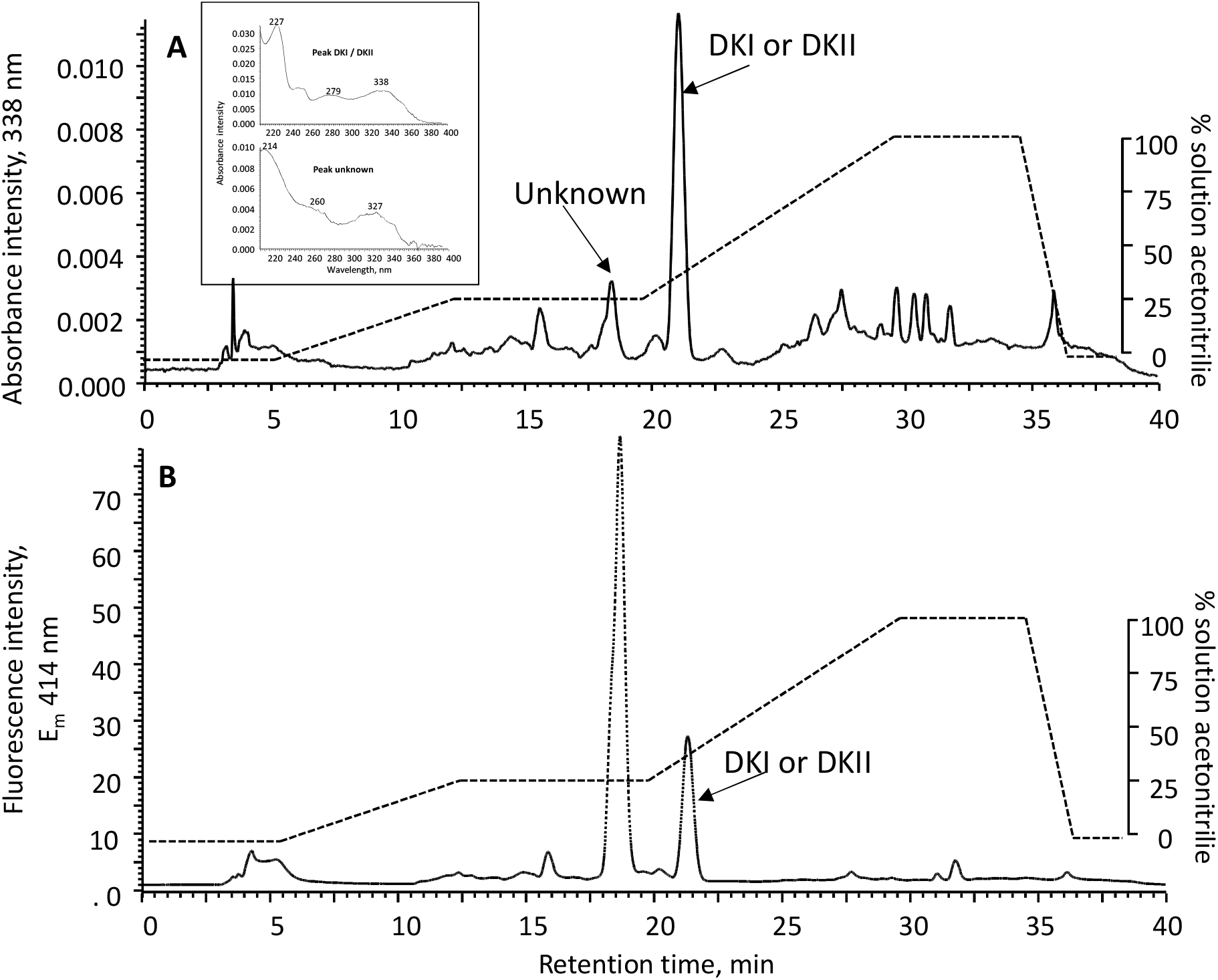
Chromatogram of the partially pure DKI or DKII samples. A) Absorbance and B) Fluorescence detections. Inserts showed spectra of the DKI or DKII-oPD, and of the minor unknown metabolite. Only, the peak eluted at 21 min, based on the similarity of the structure with KDG showed the typical spectrum as reported in Figure 4. It should be noticed that the unknown metabolite adduct is characterized by a greater yield in fluorescence emission upon excitation at 338 nm.

### Linearity

Linearity of the method was confirmed by preparing KDG standard curves for the analytical range of 0-5 nanomoles (Figure 6). Statistical analysis using Least Square Regression (LSR) indicated excellent linearity in the mentioned range, with a coefficient of determination of R^2^ ≥ 0.993 for all standard curves with fluorescence detection and R^2^ > 0.988 for the UV detection. The slopes of the calibration curves were 0.0026 and 6.64 for absorbance and fluorescence intensity detection, respectively (Figure 6A, 6B). From the respective slopes, the fluorescence intensity detection is more sensitive than the absorbance intensity detection, with a ratio of around 2500-fold. The method was compared to the thiobarbituric assay (Figure 6C). In the latter, the absorbance could not be measured below 200 nanomoles, far above the range of UV/Fluorescence detection. The slope was of 0.0007, making UV or fluorescence detection more sensitive, with a ratio of around 3.7- and 9500-fold respectively.

**Figure 6:**
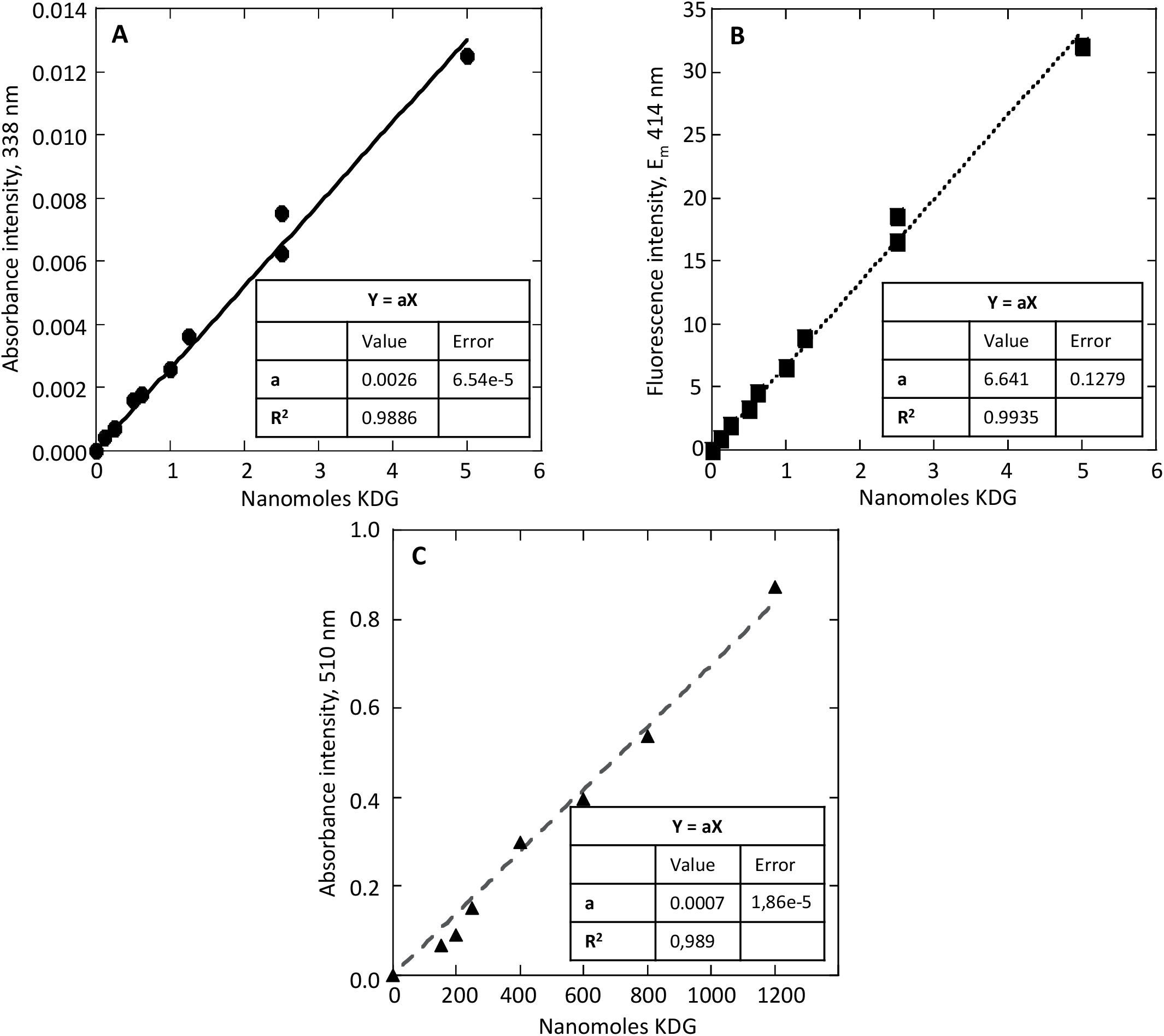
Standard curves for the pure KDG metabolite. A) Absorbance intensity at 338 nm; B) Fluorescence intensity at 414 nm upon excitation at 338 nm and C) Absorbance intensity at 510 nm in the thiobarbituric assay. Inserts reported the value of the linear slope (a) with R^2^ being the coefficient of determination. As noticed, the fluorescence intensity is more accurate than the absorbance intensity detection in our experimental set-up of the fluorimeter (2500-fold more sensitive, see y-axis), and is also more accurate than the thiobarbituric assay in terms of amounts of compounds needed for detection (x-axis).

### Limit of Detection and Limit of Quantification

The range of application of the method was verified by analyzing replicate samples containing the metabolite at various concentrations. The Limit of Detection (LOD) and Limit of Quantification (LOQ) were calculated by the linear regression method: the LOD is given by 3S_a_/b and LOQ is given by 10S_a_/b, where S_a_ is the standard error of the intercept and b is the slope of the calibration curve [46]. Based on these equations for absorbance and fluorescence detection (Figure 6), the calculated LOD values were found to be 0.093 nanomoles and 0.077 nanomoles respectively, and LOQ values were found to be 0.28 nanomoles and 0.23 nanomoles respectively, in our experimental condition settings. Compared to previous HPLC methods [34], showing a sensitivity limit of around 20 micromoles KDG, our fluorescence method detects KDG in the nanomole/femtomole range (Figure 6).

### Biological Sample Analysis

In order to test the method for KDG quantification, we used the *Dickeya dadantii* wild type strain 3937 and its *kduI*, *kduD* and *kdgK* mutant derivatives. Bacteria were grown on a glucose minimal medium supplemented or not with polygalacturonate (PGA), and samples were collected after 10 hours of growth, corresponding to the transition to stationary phase. The characteristic peak of KDG appeared only in the presence of addition of PGA in the assay, and was absent in cultures performed in glucose, as expected. Figure 7 depicts the full complexity of the chromatogram for the wild type bacterial samples with and without PGA. Meanwhile, KDG is eluted at around 16 min in a symmetrical peak. First of all, amplification of the resolution of the gradient in the isocratic phase (Figure 8) showed the peak of KDG. Secondly, the nature of the product is confirmed by coelution with pure KDG added in the assay (Figure 9). Thirdly, when working with the mutant strain *kdgK*, a major peak is amplified at 16 min (Figure 10). Additionally, chromatogram of the mutant strains *kduI* and *kduD* recovered one peak with similar spectrum signature at 21 min, plus KDG at 16 min probably resulting from the hexuronate pathway (Figure 11 and 12).

**Figure 7:**
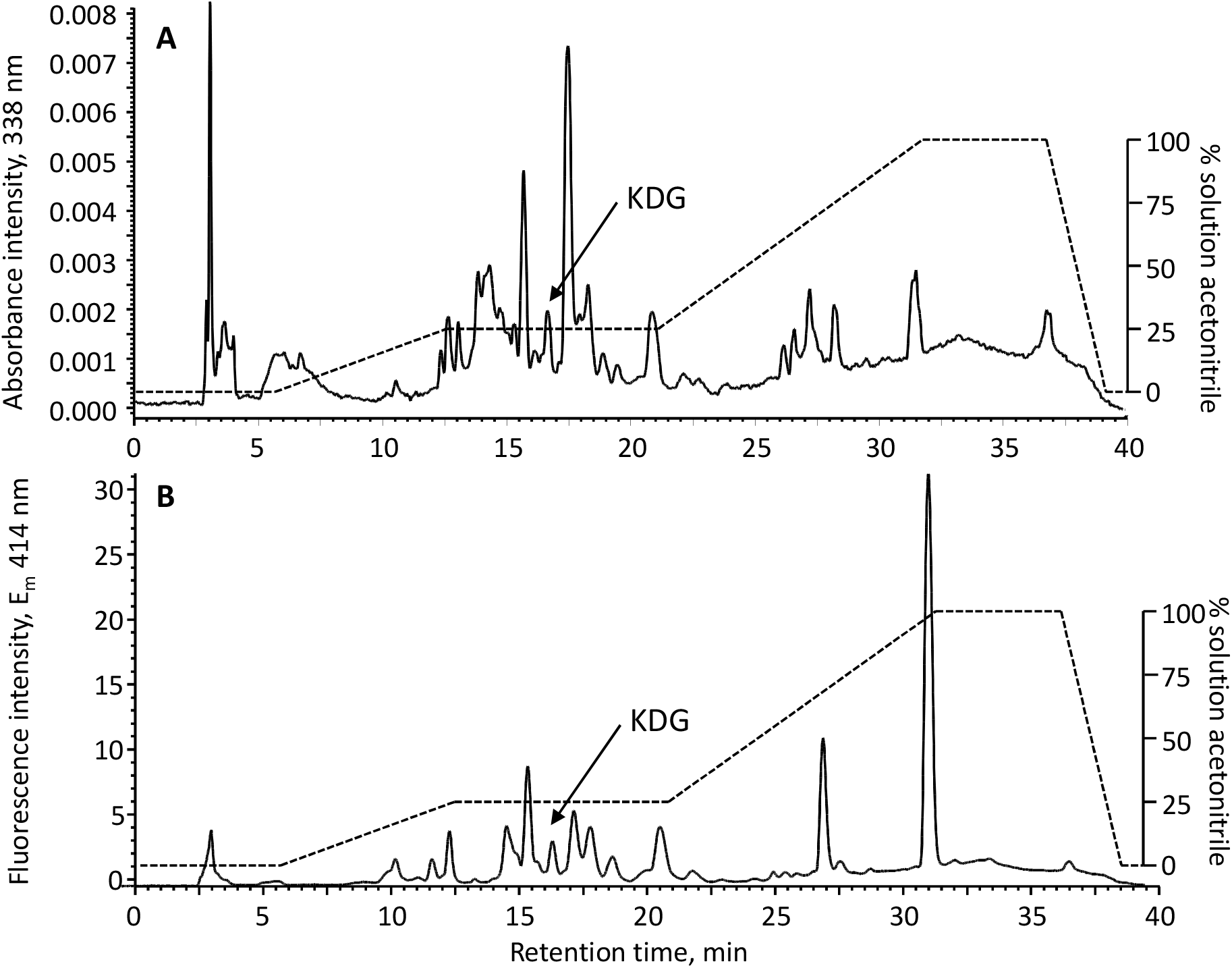
Full chromatogram of sample from wild type bacterial cells grown in glucose medium in presence of polygalacturonate (PGA). The figure highlights the complexity of the profile with crude extracts when looking to the absorbance intensity at 338 nm (A) and the fluorescence detection (B). Nevertheless, KDG peak was detected, and was clearly identified from its spectral signature as reported in Figure 4.

**Figure 8:**
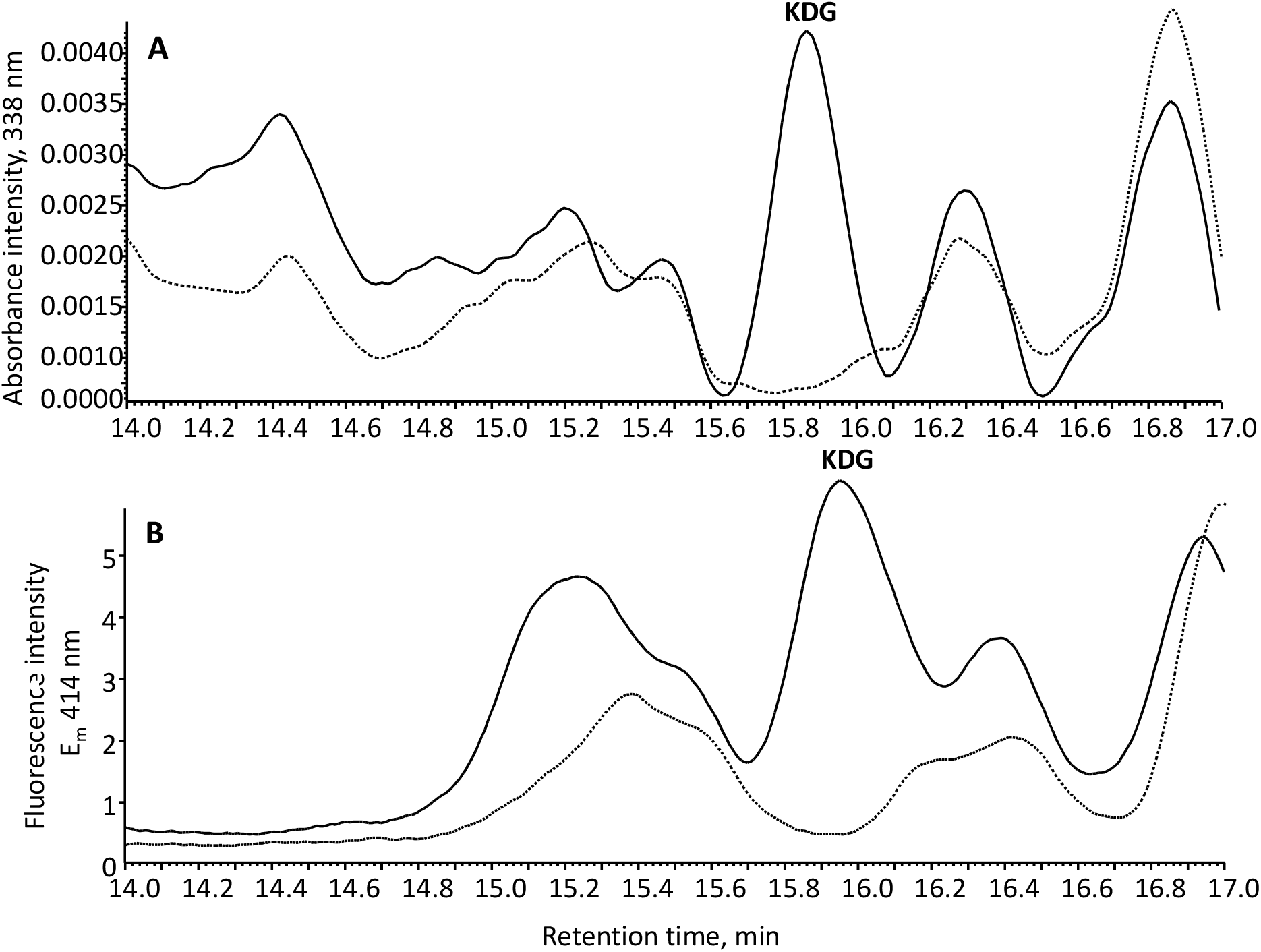
Wild type profile chromatogram. Sample from wild-type bacterial cells grown in glucose medium in the presence (continuous black line) and absence (gray dotted line) of polygalacturonate (PGA) were plotted. The chromatogram is in the isocratic phase of the gradient to amplify the image for resolution of KDG-oPD adduct molecule. A) Absorbance intensity at 338 nm and B) Fluorescence intensity at 414 nm.

**Figure 9:**
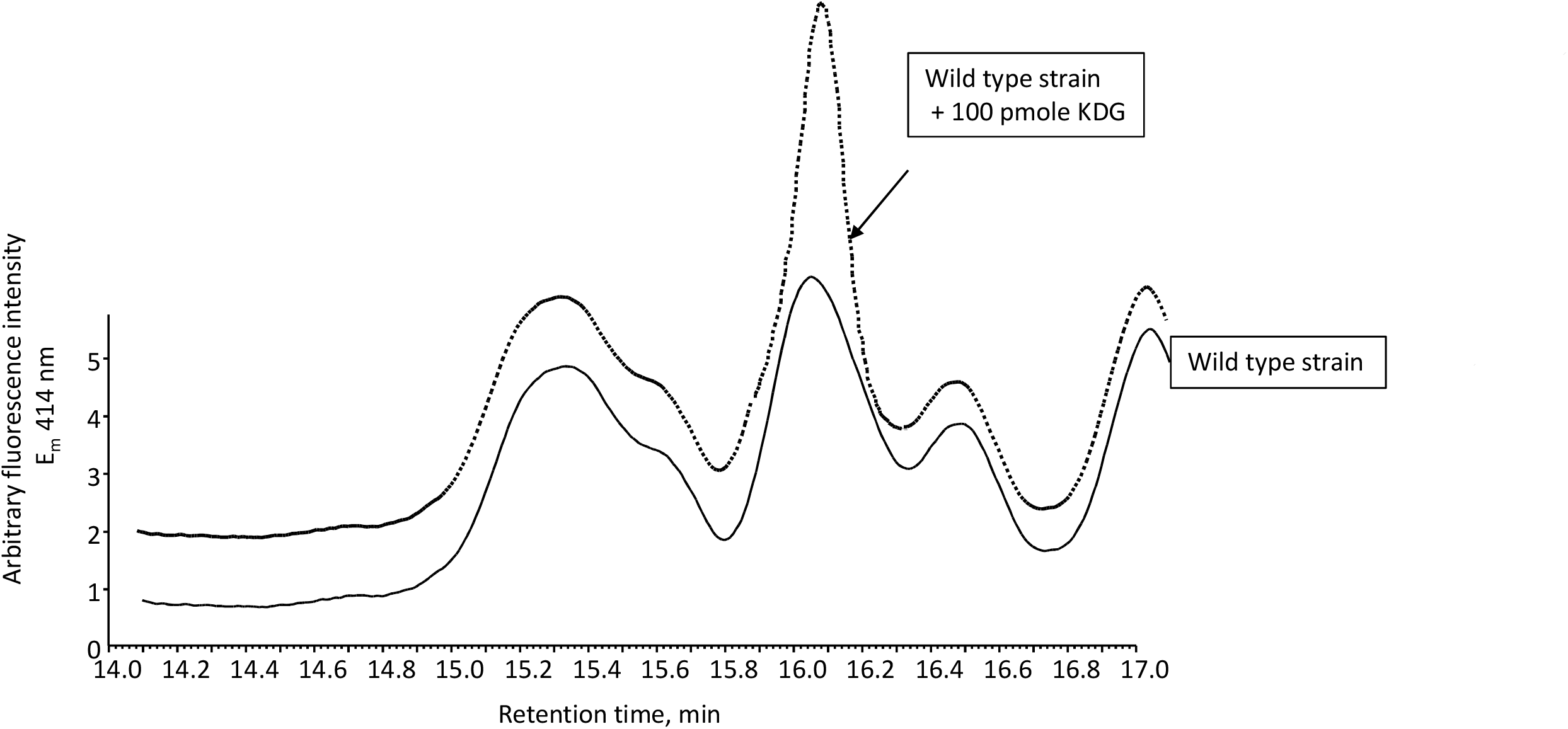
Coelution chromatogram of sample from wild-type bacterial cells. The cells were grown in glucose medium in the presence of polygalaturonate (continuous black line) and with 100 pmoles of commercial KDG (gray dotted lines). The chromatogram is in the isocratic phase of the gradient to amplify the image for resolution of KDG-oPD adduct molecule.

**Figure 10:**
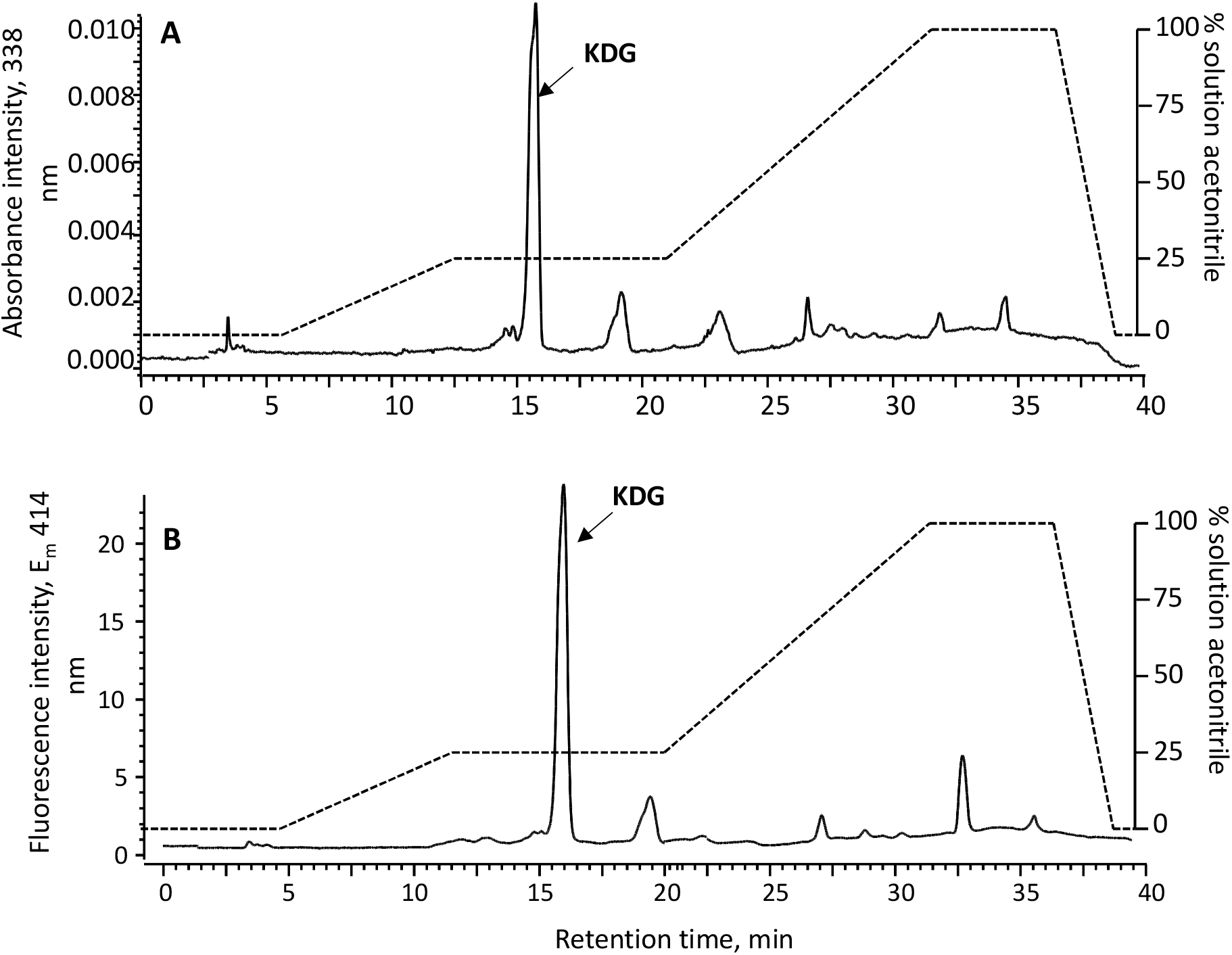
Chromatogram of *kdgK* mutant strain of *Dickeya dadantii*. The cells were grown in glucose medium in presence of presence of polygalacturonate (PGA). A) Absorbance intensity at 338 nm and B) Fluorescence intensity at 414 nm. It should be noticed the difference in the y-scale in each chromatogram indicating a strongest sensitivity in the fluorescence over the absorbance detection. Furthermore, this chromatogram is a further support for the elution of KDG at 16 min as reported in Figure 3, 7, 8 and 9 in the co-elution experiment. Moreover, the peak of KDG is the sum resulting from catabolism of the saturated and the unsaturated digalacturonate (see Figure 1).

**Figure 11:**
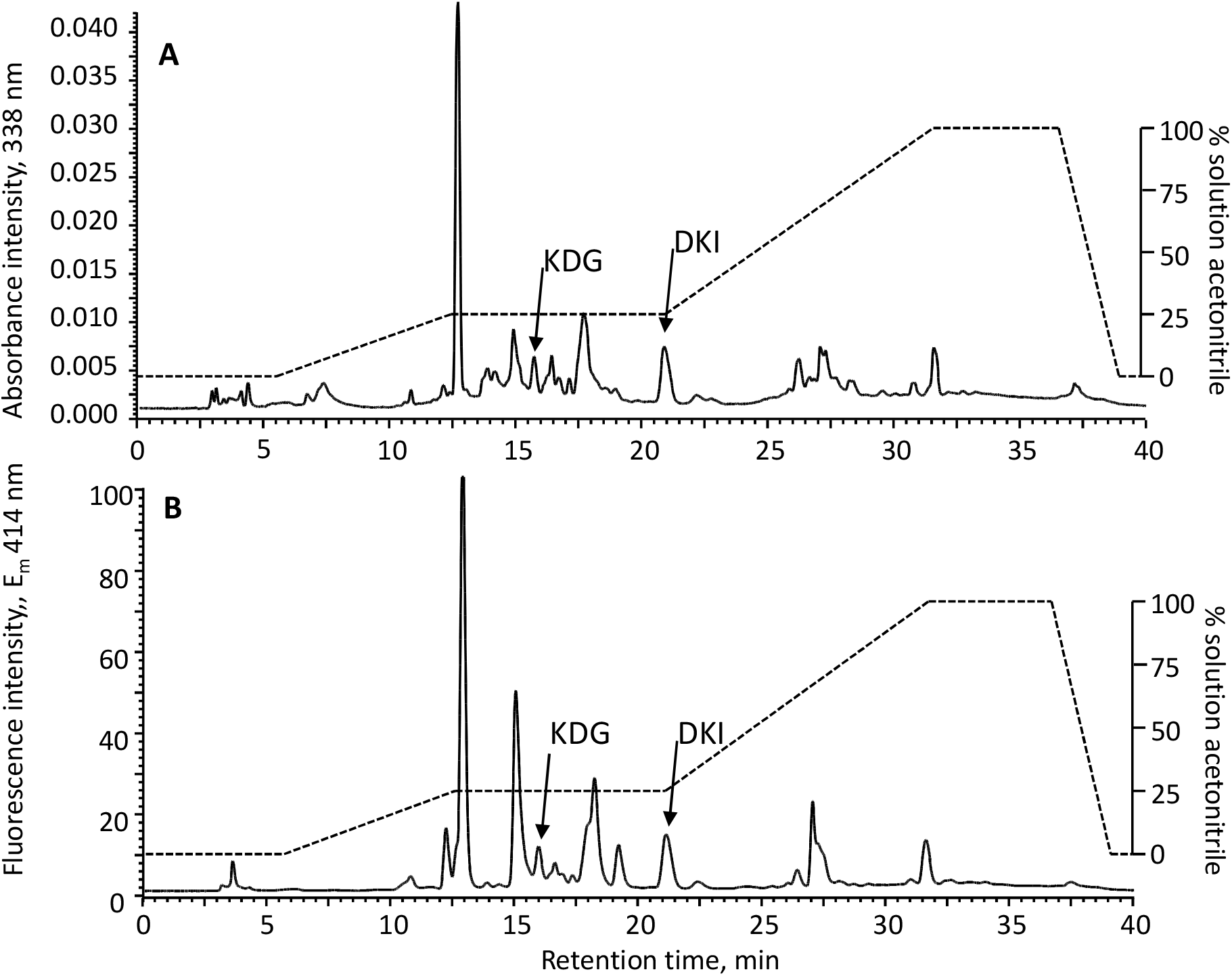
Chromatogram of *kduI* mutant strain of *Dickeya dadantii*. Cells were grown in glucose medium in presence of presence of polygalacturonate (PGA). A) Absorbance intensity at 338 nm and B) Fluorescence intensity at 414 nm. It should be noticed the difference in the y-scale in each chromatogram indicating a strongest sensitivity in the fluorescence over the absorbance detection. Furthermore, DKI / DKII eluted at around 21 min as observed in Figure 5 with the partially purified compound. Moreover, a pic of KDG is detected in the mutant, originating from the saturated digalacturonate viathe parallel hexuronate pathway.

**Figure 12:**
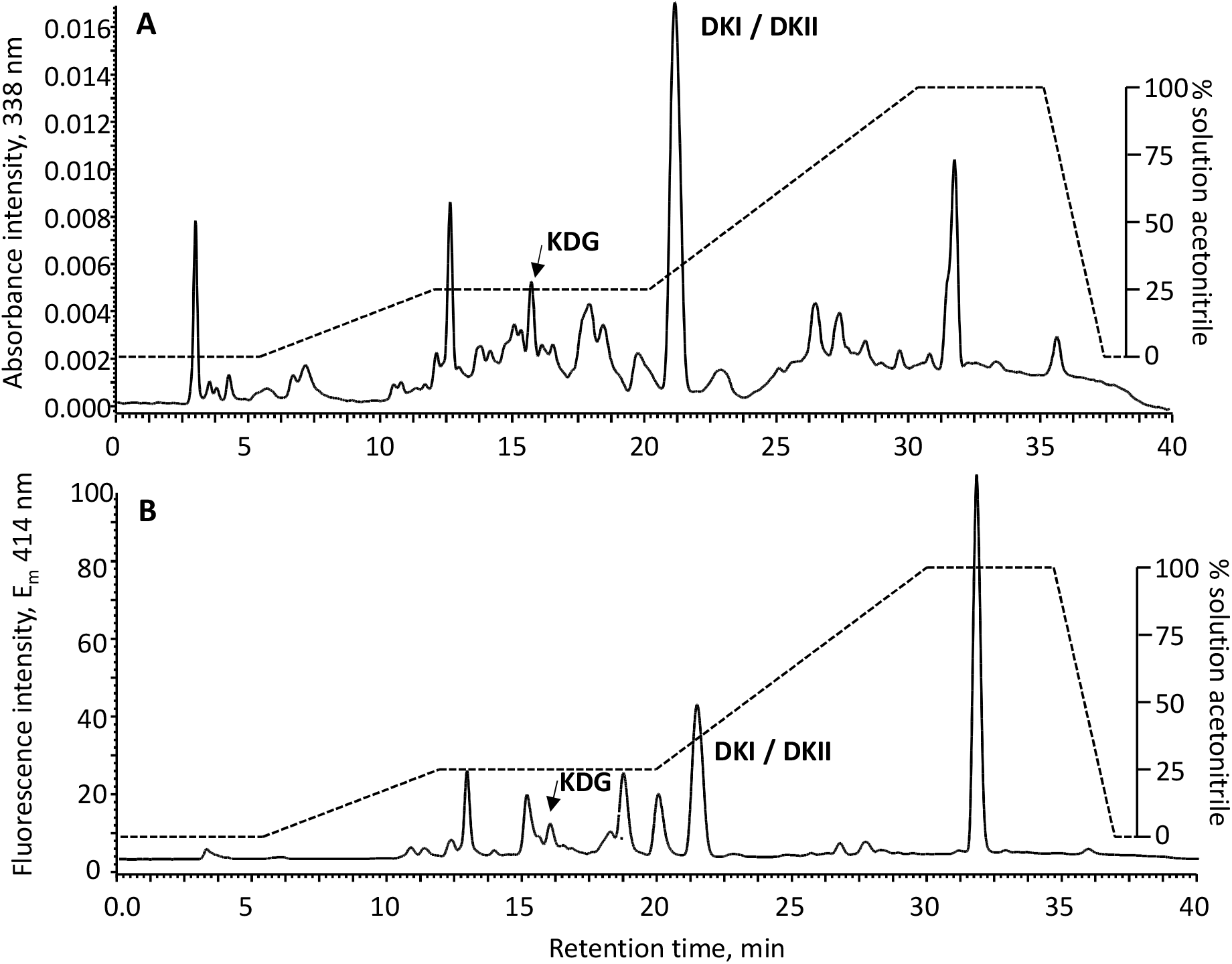
Chromatogram of *kduD* mutant strain of *Dickeya dadantii*. The cells were grown in glucose medium in presence of presence of polygalacturonate (PGA). A) Absorbance intensity at 338 nm and B) Fluorescence intensity at 414 nm. It should be noticed the difference in the y-scale in each chromatogram indicating a strongest sensitivity in the fluorescence over the absorbance detection. Furthermore, DKI / DKII eluted at around 21 min as observed in Figure 5 with the partially purified compound. Moreover, a pic of KDG is detected in the mutant, originating from the saturated digalacturonate via the parallel hexuronate pathway.

### Quantitation of metabolites

The KDG concentration was quantified in the wild type *D. dadantii* strain and the three mutant derivatives (*kduI*, *kduD* and *kdgK*) to study the formation of intracellular concentration of KDG (Figure 13). The intracellular quantification was carried out based on the assumption of a cell volume to be 3.6 μL.mL^−1^.OD^−1^ where optical density (OD) is measured at 600 nm [47]. In the medium supplemented with a PGA source the maximum quantification of KDG is observed in the *kdgK* mutant, as expected since it accumulates the KDG metabolite (up to 233 mM). The *kduI* and *kduD* mutants accumulated KDG at higher level than wild type cells (10- to 30-times respectively), due to the increased catabolism of saturated digalacturonate in the hexuronate pathway. The Pel enzymatic activities assayed in the mutants also are consistent with the elevated levels of KDG (Table 2).

**Table 2:** Pel activity in WT, *kdgK*, *kduI*, *kduD* mutants Pectate lyase production of *Dickeya dadantii* wild type strain and mutant strains grown in minimal medium supplemented with glucose or glucose + polygalacturonate (PGA). Samples were taken in late stationary phase (10 hours in our conditions). Pel specific activity is expressed as μmol of unsaturated product liberated per min per mg of bacterial dry weight (n = 4, and values are means +/− standard errors).

| Strain | Growth medium | Pel activity |
| --- | --- | --- |
| WT | Glucose | 0.45 +/- 0.25 |
|  | Glucose + PGA | 11.30 +/- 3.41 |
| *kduI* | Glucose | 0.42 +/- 0.24 |
|  | Glucose + PGA | 11.40 +/- 4.91 |
| *kduD* | Glucose | 0.41 +/- 0.23 |
|  | Glucose + PGA | 12.27 +/- 4.57 |
| *kdgK* | Glucose | 0.54 +/- 0.16 |
|  | Glucose + PGA | 36.45 +/- 9.67 |

**Figure 13:**
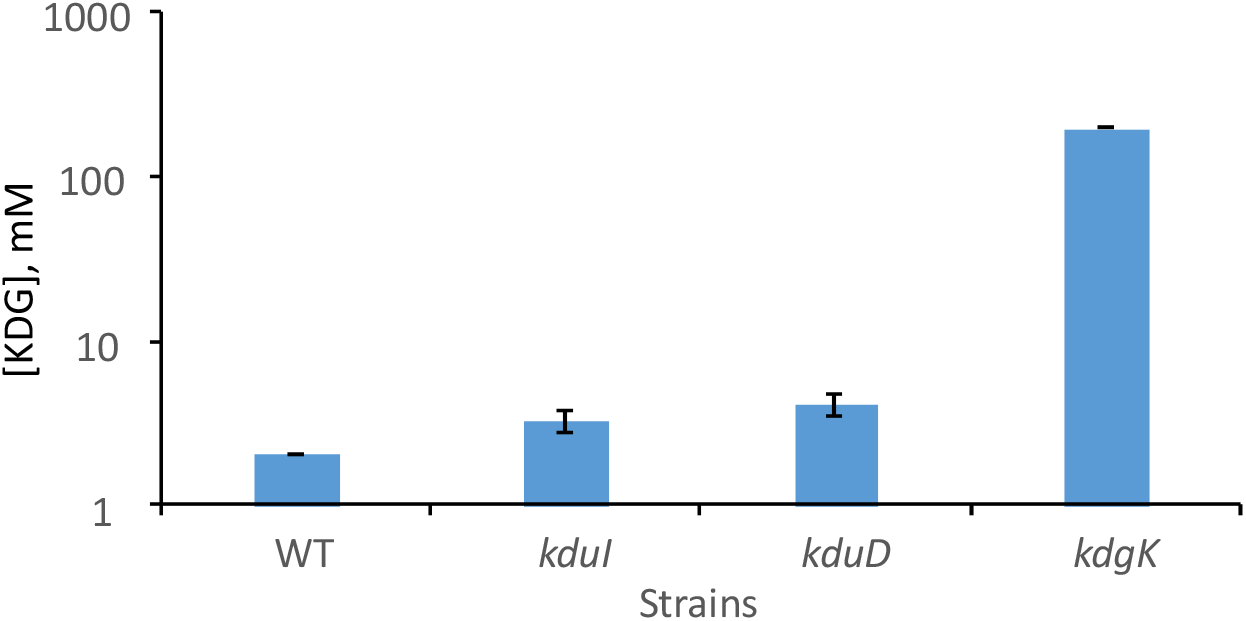
Logarithmic representation of KDG concentrations. The concentrations were measured in wild type *Dickeya dadantii* and *kduI*, *kduD*, *kdgK* mutants affected from steps of the pathway of unsaturated digalacturonate (Figure 1). Error bars indicate the standard error observed in experimental quantifications over independent biological triplicates (n = 3).

DKI and DKII could not be further differentiated on the chromatogram due to their structural similarity and unavailability of pure DKI and DKII compounds. We quantified DKI and DKII based on the KDG calibration curve considering that the absorbance and fluorescence slopes would be similar. In *kduI* mutant, DKI was not converted into DKII, and its concentration was quantified around 15mM. In the *kduD* mutant the combined concentration of DKI+DKII was 31mM. The *kdgK* mutant had no detectable DKI or DKII molecules.

## Conclusion

To demonstrate that our technique is specific for KDG alone, we showed that the retention time for the derivatives DKI/DKII is 21 minutes whereas the retention time for KDG is around 16 minutes. Nonetheless, this method is solid enough to eliminate the suspicion of cross derivatization between KDG and DKI/DKII. We do not consider galacturonate pathway since the similarity in the structure of compounds are not significant for the possibility of cross derivatization.

The fluorescent technique was 2500-fold more sensitive over the absorbance detection. The results also indicate the higher reliability of the fluorescence technique over the absorbance detection method since it has a lower coefficient of variation. Nonetheless, the combination of the two techniques provide further validation of the results and stronger quantification.

Thus, we describe a robust method for the quantification of the metabolite KDG from biological samples. This method will help in the analysis and quantitative modelling of gene regulatory networks where KDG is known to have any effect [7,28]. Moreover, quantifying metabolite KDG flux is critical for biofuel production during pectin and alginate degradation or for other high value products derived from these two polymers. There is also potential scope for the industrial application of this technique in metabolic engineering of compounds like Vitamin C from KDG [48].

## Acknowledgements

Authors are grateful to G Condemine and N Hugouvieux-Cotte-Pattat for providing the DKI/DKII samples, and for discussion and critical readings.

